# Mechanisms of ATM Inhibitor AZD1390-Mediated Radiosensitization by Comparing DNA DSB Formation and Repair in 4T1 Cells

**DOI:** 10.1101/2025.07.27.667090

**Authors:** Jake Atkinson, Joshua Chopin, Eva Bezak, Hien Le, Ivan Kempson

## Abstract

Radioresistant cancers often exhibit upregulated DNA damage response (DDR) proteins including ataxia telangiectasia mutated (ATM), recovering more readily from radiation-induced DNA double-strand breaks (DSBs). ATM inhibitors (ATMi) are therefore being explored as adjuvants in radiotherapy to both enhance radiosensitivity and minimise normal tissue complications. The molecule AZD1390 has been developed as an inhibitor of ATM and is undergoing assessment in clinical trials. However, traditional markers of DNA DSB repair, such as γH2AX, represent stages of DDR downstream from the action of ATM. The ATMi AZD1390, developed by AstraZeneca, ultimately prevents phosphorylation of the H2AX histone variant. In order to quantify ATMi AZD1390 action, the novel and highly complementary assay SensiTive Recognition of Individual DNA Ends (STRIDE) was employed to directly quantify DSBs in comparison to γH2AX after X-ray irradiation of 4T1 cells. Findings revealed that ATM inhibition via AZD1390 delays DSB repair initiation and appears to play a greater role in suppressing DDR beyond ATM inhibition alone. In X-ray irradiated conditions, obfuscation of DSBs commenced between 30- to 45-minutes post-irradiation without AZD1390 versus 45 to 60-minutes for cells pretreated with AZD1390 and entirely prevented γH2AX in the majority of cells. STRIDE and γH2AX exhibited almost no co-localization indicating that they provide distinct and complementary information. DSB formation was also assessed in cells fixed pre-insult to minimise any biological response, providing unprecedented assessment of DSB formation. These results highlight the potential of STRIDE to accurately measure DNA damage response kinetics, paving the way for more precise mechanistic studies into the role of ATM in radiotherapy.

## Introduction

During X-ray radiotherapy, one of the principal modes of cell death is the induction of DNA double-stranded breaks (DSBs). These occur through either direct ionization of DNA or indirectly via ionization of surrounding water molecules, generating cytotoxic reactive oxygen species (ROS) which subsequently interact with DNA. If unrepaired, DSBs lead to mutations, chromosomal aberrations, or cell death, making efficient detection and repair critical for maintaining genomic integrity [1]. As such, cells constantly monitor DNA damage through activation of key signalling molecules within the DNA damage response (DDR). This includes ATM (ataxia telangiectasia mutated), ATR (ATM and Rad3-related), and γH2AX, which enable the recruitment of factors that initiate DSB repair pathways. ATM plays an intricate role in regulating the DDR, being recruited to DNA breakage sites by the DSB tethering MRE11-RAD50-NBS1 (MRN) complex, and subsequently undergoing autophosphorylation at Ser1981 [2]. This active form of ATM is then able to phosphorylate Ser139 of histone variant H2AX (γH2AX), which allows for widespread chromatin remodelling and accumulation of proteins within the homologous recombination (HR) or non-homologous end joining (NHEJ) pathways at DSB sites [3].

In clinical situations, the effectiveness of single modality radiation or chemotherapy is often constrained by the risk of normal tissue complications. Therefore, a combination of surgery, radiotherapy, and chemotherapy is often prescribed in advanced cancers to enhance the likelihood of tumour control and eliminate microscopic disease [4]. Dysregulation of the DDR often leads to radioresistance, making proteins within this pathway candidates for targeted chemotherapies [5]. Development of ATM inhibitors (ATMi) therefore holds great potential in increasing the vulnerability of radio- and chemo-resistant tumours. Inhibiting ATM can enhance the sensitivity of cancer cells to radiotherapy by interrupting the DDR and preventing cancerous cells from surviving otherwise lethal damage [6, 7]. AZD1390, developed by AstraZeneca, is one such inhibitor highly selective against ATM with a half-maximal inhibitory concentration (IC50) of 0.78 nM [8]. It possesses an advantage over similar ATMi, including the previously developed AZD0156, due to its ability to penetrate the blood-brain barrier (BBB). The BBB prevents many widely used anti-cancer pharmaceuticals from entering the brain [9], and even if penetrance is possible, intratumoural drug concentration may not exceed the cytotoxic threshold required to eliminate the tumour entirely [10]. Therefore, the combination of both AZD1390’s selectivity and BBB penetrance is highly desirable for treatment of radioresistant brain tumours such as glioblastoma multiforme (GBM) [9, 11-13]. This capability was validated in a study using 11C-labelled AZD1390 and positron emission topography (PET) imaging in cynomolgus monkey brains [8], and a phase I clinical trial (ClinicalTrials.gov ID: NCT03423628) has also commenced to determine the tolerability of administering AZD1390 in combination with radiotherapy in GBM patients [9].

ATMi have also been investigated for their role in enhancing the radiosensitivity of breast cancers (BC). As the most commonly diagnosed cancer in women [14], BC commonly shows abnormalities in the ATM gene, which is a contributing risk factor in both its development [15, 16] and cancer-specific survival [17]. Inhibition of ATM has been shown to reduce the degree of cell invasion, migration and metastasis in both MDA-MB-231 and MCF-7 cells [18]. AZD1390 has been tested in mice against patient derived xenografts of HER2+ and triple-negative breast cancer, with pre-treatment sensitizing these cancer types to radiation in orthotopic models [19]. AZD1390-induced ATMi has also been shown to delay growth in 4T1 tumour models, and activate the cGAS/STING pathway, enhancing the type 1 interferon response and increasing the effectiveness of anti-PD-1 immunotherapy [20]. These findings highlight the potential of ATM inhibitors to broaden the therapeutic window and improve adjuvant therapies in BC treatment.

There is a need for more sensitive and precise techniques to assess DNA damage and repair, particularly within the realm of drug development and mechanistic studies. A major limitation in the characterization of ATMi function currently is the inability to accurately quantify their impact on DNA damage and repair. Conventional immunofluorescence-based methods typically rely on the detection of DNA repair markers, yet these assays do not directly quantify DSBs [21] and are prone to biases [22, 23]. For instance, using markers such as ATM or γH2AX, which signal downstream DNA damage response molecules to initiate repair, becomes problematic when DDR inhibitors are applied. Inhibition of ATM leads to reduced phosphorylation of γH2AX [24], which diminishes its detection via immunofluorescence and, as a result, compromises the reliability of these markers as indicators of DNA damage. γH2AX is the most conventional proxy for DSBs, however this signal is not expressed when ATMis are used [21]. Therefore, no published studies have been able to describe the exact impact of ATMi on the formation and repair of DSBs by the DDR with immunofluorescent approaches. The effectiveness of ATMis in modulating DSB repair cannot be fully understood using current assays, impeding efforts to assess their role in enhancing genotoxicity.

There are significant gaps in understanding how ATMis affect DSB repair kinetics. Novel immunofluorescence assay STRIDE (SensiTive Recognition of Individual DNA Ends) offers a highly sensitive approach to address this challenge by detecting DSBs at single-break resolution [25]. This enables a precise analysis of the effects of ATM inhibition on DNA repair processes, particularly in the context of radiation-induced DNA damage. As STRIDE can bind to free DSB ends [25], this allows the ability to quantify the physical DNA breakages caused by X-rays and provide a comparison to biological signals activated in response. By leveraging the sensitivity of STRIDE, the primary aim of this study was to characterize the role AZD1390 plays in influencing the repair of DSBs. Internalization of AZD1390 was anticipated to inhibit activity of γH2AX, and consequently perturb DSB repair. Immunofluorescence stains for both STRIDE and γH2AX were used to quantify DSB formation and repair in 4T1 cells, a murine model of triple-negative BC, exposed to ATM inhibitor AZD1390, X-rays, and their combination. Capturing these changes in DSB and repair dynamics presents a powerful tool to provide insight into how DDR inhibitors modulate the DNA damage repair landscape, especially in the context of radiosensitization strategies. This study presents a proof-of-concept protocol developed to evaluate DNA damage and repair mechanisms in the context of DDR inhibitors such as AZD1390, providing insights into the impact of ATMis on DNA repair processes. While the experiments conducted using 4T1 cells provide a model for these mechanisms, the results may not fully represent the complexities inherent in different normal and malignant cell types. These findings establish a foundation for further investigations into the broader applicability of DDR inhibitors in diverse cellular contexts, with the aim of better informing drug development.

## Methods

### Cell Culture and Pre-treatment

Murine 4T1 cells (ATCC, CRL-2539) were maintained in RPMI medium (Thermo Fisher, 11875093) incubated at 37°C, 95% relative humidity and 5% CO^²^. 12 hours before irradiation, cells were seeded into Ibidi µ-Slide 8 Well chambered coverslips (Ibidi, 80826) at a density of 40,000 cells per well. To assess the impact of ATM inhibition, AZD1390 (MedChemExpress, HY-109566) was introduced at a final concentration of 30 nM to designated wells two hours before irradiation. All conditions were performed in duplicate.

### X-ray Irradiation

The x-ray source was the 160 kV RadSource 2000 Small Animal Irradiator at the South Australian Health and Medical Research Institute (SAHMRI) in Adelaide, Australia. Cells were irradiated for 45 seconds in 200 µL culture medium, delivering a total dose measured to be approximately 1.93 Gy (0.0429 Gy/s) determined by a Radcal Accu-Dose+ probe positioned adjacent to the slides in air. The dose received by the cells varies depending on the back-scattering material, volume of PBS above the cells, and surrounding scattering materials. The 1.93 Gy recorded from the probe serves as an estimated dose received by the cells but expect this to be within 10% of the actual dose. Fixation was performed prior to irradiation (0-minute timepoint), and at multiple post-irradiation (post-IR) intervals: 15, 30, 45, 60, 120, 240, and 1440 minutes. Cells were fixed using 4% paraformaldehyde for 10 minutes at room temperature.

### STRIDE Assay

After fixation, cells were permeabilized using PBS containing 0.5% Triton X-100 for 10 minutes, followed by a 10-minute incubation with 0.5–10 mM EDTA/PBS at room temperature. Samples were subsequently rinsed with 100 µL of TUNEL kit Wash Buffer. DNA DSBs were labelled with BrdUTP using the APO-BrdU Kit (BD Biosciences, Cat No. 556405), in accordance with the manufacturer’s protocol. A master mix containing BrdUTP, TdT enzyme, TdT reaction buffer, and ddH^²^O (50 µL per well) was applied and incubated at for 1 hour at 37°C. Wells were then rinsed with 100 µL of Rinse Buffer and blocked with 3% BSA (200 µL per well) for 1 hour at 37°C. Primary antibodies, Mouse MoBu-1 anti-BrdU (Abcam, 8039, 1:500) and Rabbit Polyclonal anti-BrdU (Abcam, ab152095, 1:200) were incubated overnight at 4°C in 3% BSA. Wells were rinsed again with Duolink® In Situ Wash Buffers, followed by addition of two secondary antibodies: Duolink® PLA® Probes Anti-Rabbit PLUS (Sigma Aldrich, DUO92002) and Anti-Mouse MINUS (Sigma Aldrich, DUO92004) were added for 1 hour at 37°C. This was followed by ligation for 30 minutes at 37°C and polymerase amplification for 100 minutes, with washes between steps. Cells were then finally washed with 1X and 0.01X Wash Buffer B.

### γH2AX Assay

Blocking was performed with 3% BSA for 1 hour at 37°C before incubation with anti-phospho-H2A.X (Ser139) Rabbit mAb conjugated with Alexa Fluor 488 (Cell Signaling Technology, #9719). 1 mg/mL DAPI (Thermo Fisher, Cat No. 62248) was used for nuclear staining.

### Confocal Imaging

Immunofluorescent images were captured with the Zeiss LSM 710 confocal microscope (Carl Zeiss, Germany) with a Plan-Apochromat 20x/0.8 M27 lens. For each condition, four fields of view, each comprising approximately 24 z-slices (2 µm) with 10% overlap, were acquired. The final stitched images were 1950 × 1950 pixels with a size of x: 807.68 µm, y: 807.68 µm, z: 48.00 µm. Excitation and emission spectra were: STRIDE (λex 644 nm; λem 669 nm), γH2AX (λex 488 nm; λem 496 nm), DAPI (λex 350 nm; λem 470 nm).

### Image Analysis

Confocal microscopy images were assembled into stitched composites, and z-stack maximum projections were generated using ZEN (black edition) 3.0 SR before being exported in .TIF format. These maximum projection images were subsequently analysed in CellProfiler 4.2.6 [26] which provides modular tools for batch processing large image datasets. The software’s workflow is adaptable, allowing modules to be added, removed, or rearranged as required by each specific experimental setup. Object identification within nuclear regions was performed by decomposing RGB images into their grayscale components, detecting and segmenting nuclei using the DAPI signal (blue channel), and then quantifying fluorescence intensity from the red, green, and blue channels within each nucleus. For high-magnification (63x) images, thresholding techniques were applied to generate masks of individual STRIDE and γH2AX foci, enabling per-nucleus quantification. The extracted data was exported to Excel, where STRIDE and γH2AX intensity values were normalized against DAPI intensity before being imported into GraphPad Prism 10 for visualization and statistical evaluation. Given the non-parametric nature of the dataset, statistical significance was assessed using the Mann-Whitney U test at selected post-irradiation timepoints to compare conditions with and without AZD1390 treatment. Final figure preparation involved adding scale bars in ImageJ 1.54f and assembling confocal image collages using Photopea.

## Results and Discussion

### Confocal Imaging

Two-hours prior to irradiation (T = -120 minutes), cells were treated with or without 30 nM AZD1390 and then returned to a 37°C humidified incubator. Wells were then fixed at a 0-minutes timepoint, that is at the 2-hour exposure point for the AZD1390 pre-treatment and just prior to X-ray irradiation. The rationale was to obtain a measure of the physical damage caused by X-rays without initiation of the biological DDR. As of the time of writing and to the best of our knowledge, this is the first report of such a procedure prior to irradiation to determine DSB counts *in vitro*. After irradiation, cells were also fixed incrementally from 15 to 1440 minutes post-insult to compare both STRIDE and γH2AX kinetics in single cells. Confocal imaging was performed at 20x to obtain integrated intensity values for all experimental conditions and timepoints, and at 63x for foci quantification of the 15-, 30- and 45-mintutes post-IR ‘X-ray irradiation only’ conditions (Figure 1, Figure 6).

**Figure 1:**
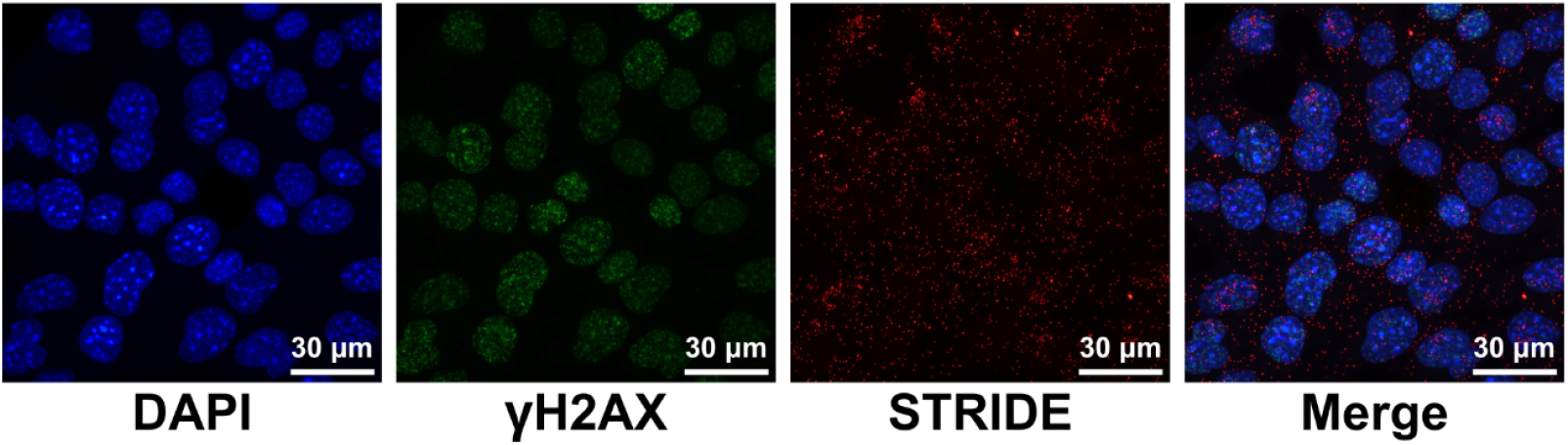
Confocal images at 63x magnification of 4T1 cells irradiated with 1.93 Gy of 160 keV X-rays and fixed 30-minutes post-IR. Cells were co-stained to identify DSBs (red) and γH2AX (green) with STRIDE (Duolink In Situ Detection Reagents FarRed (Cy5)), and anti-phospho-histone H2AX conjugated with Alexa 488 respectively. DAPI was used to label individual nuclei.

### 0 Gy X-rays Negative Controls

For the negative control with no radiation and no ATMi (Figure 2), little to no STRIDE foci were expected to form as well as a relatively low, baseline level of H2AX phosphorylation across all timepoints. This is because STRIDE foci, representative of DSBs, should mainly occur due to X-ray irradiation of cells. As there was no irradiation of these samples, STRIDE fluorescence should not be abundant under this condition. Generally, the majority of conditions represent values from between ∼700 - ∼1,500 individual cells (**Error! Reference source not found**.). Data are also presented in terms of the mean values in Figure S1, which is useful for easier visualization of the data trends. Potential sources of STRIDE foci formation in this negative control include spontaneous DSBs during DNA replication, cytosolic DNA fragments or damaged mitochondrial DNA. Immunofluorescence imaging of these control samples does not suggest that these sources of STRIDE foci exist in high numbers, however. Since cells were not cell phase synchronized, the DNA content (and hence DAPI signal) between cells in different phases would vary. STRIDE and γH2AX signals could also present artefacts due to this, as differences in DNA content may correlate with additional DSBs and H2AX phosphorylation events. To minimize this, the integrated intensities of STRIDE and γH2AX were divided by the integrated intensity of DAPI to express these signals as a proportion of DAPI per cell.

**Figure 2:**
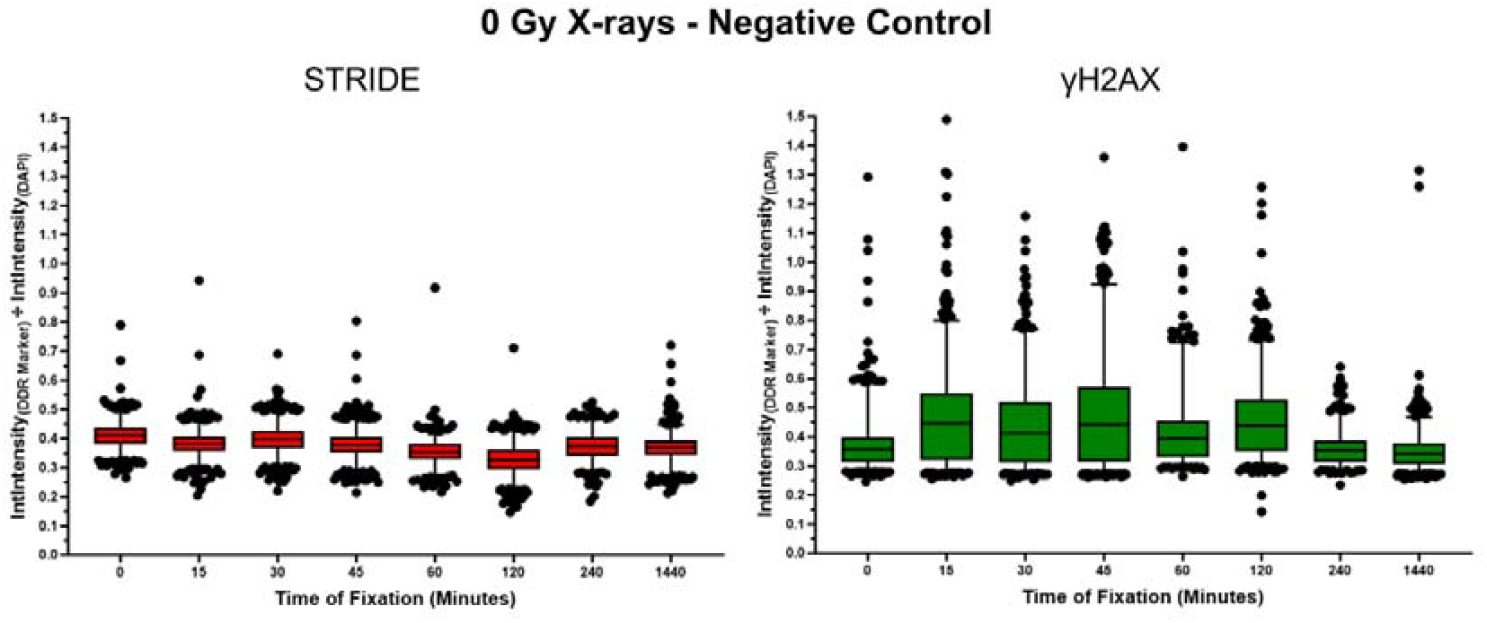
Negative control condition of 4T1 cells without X-ray or ATM inhibitor AZD1390 treatment (negative control). Integrated intensity values for STRIDE, γH2AX, and DAPI from single cells were measured. STRIDE and γH2AX values were normalised to DAPI. Data show the median value within the 25^th^ and 75^th^ percentile data points (box). Whiskers show the 2.5 - 97.5 percentile, and dots represent data points outside the range represented by whiskers.

When observing normalized integrated intensity values, both STRIDE and γH2AX reflected a low level of genotoxic stress within these cell populations. For the sake of this experiment, it can be considered the ‘baseline’ level of damage and repair respectively for 4T1 cells. Median intensities range from 0.32 - 0.41 for STRIDE and 0.34 – 0.45 for γH2AX, with the latter marker’s signal appearing to increase slightly between the 0- (0.36) and 15-minute (0.45) timepoints and then plateaus at the majority of timepoints afterwards. This could be due to the minute level of stress these cells were placed under on the trip to and from the irradiator lab. Measures were taken to ensure cells were temperature controlled by using heat packs surrounding the culture plates in their transport container, although it did not permit for maintenance of optimal oxygen and carbon dioxide levels (∼20% and 5% respectively). Additionally, as the containers the well plates were carried in were not sterile, and due to removing the lids from the Ibidi 8-well plates prior to irradiation to minimise attenuation, there was a potential window for infection although it is anticipated that over these time frames there would be negligible impact.

Data from cells which were cultured with the ATMi are provided in Figure 3. Like the negative control, there is minimal change in STRIDE signal across all timepoints. As ATM is an activator of γH2AX, inhibition of ATM through AZD1390 also results in a reduction in phosphorylation of γH2AX. AZD0156, another clinical-grade ATM inhibitor from the same family as AZD1390, was also established to suppress γH2AX in FaDu cells at a dose of 30 nM following 5 Gy irradiation [24]. Therefore, with AZD1390 being highly selective of ATM with an IC50 of 0.78 nM [8], and the dose utilised being far above this (30 nM), near-complete suppression of γH2AX signal was expected to be observed. Hence for this AZD1390 control, substantial changes in γH2AX fluorescence intensity across timepoints were not expected. Indeed, the median integrated intensities for this sample are lower across all timepoints (0.28 – 0.32) compared to cells without AZD1390. STRIDE foci exhibit comparable median intensities (0.24 – 0.34), compared to the negative control where there is also no X-ray treatment. These data indicate that there was still a low level of genotoxic stress to the cells in this condition, but the resulting γH2AX expression which had been observed in Figure 2, has been suppressed in Figure 3 due to the ATMi.

**Figure 3:**
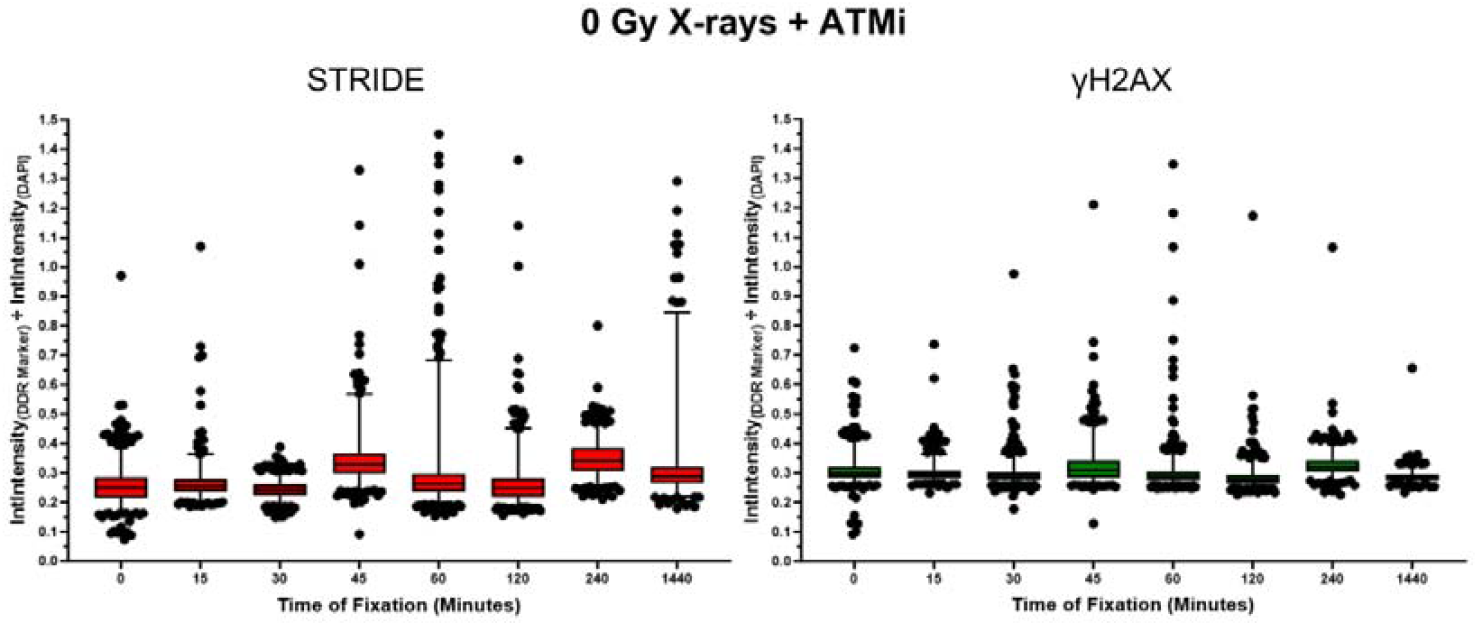
Non-irradiated 4T1 cells pre-treated with 30 nM AZD1390. Integrated intensity values for STRIDE, γH2AX, and DAPI from single cells were measured. STRIDE and γH2AX values were normalised to DAPI. Data show the median value within the 25^th^ and 75^th^ percentile data points (box). Whiskers show the 2.5 - 97.5 percentile, and dots represent data points outside the range represented by whiskers.

### X-ray Irradiation Combined with AZD1390

Upon exposure to X-rays and the induction of physical DNA lesions, the DSB kinetics of STRIDE become apparent (Figure 4). Initially, STRIDE DSB foci are at their peak at the 0-minute timepoint (0.69 arbitrary units), wherein samples were fixed pre-irradiation. This is the first time DSB’s have been measured in this way and provides a new approach to studying DNA damage in situ with no active DDR. The maximum number of DSB foci were expected to be observed at this timepoint, as the biological repair mechanisms do not yet have an opportunity to initiate repair post-IR. Therefore, it is an unambiguous representation of the exact number of DSBs that are occurring due to the physical dose administered. As time post-IR increases, STRIDE DSB intensity already decrease slightly at 15 minutes (0.59) and then maintained at 30 minutes (0.59) post-irradiation, after which there is a substantial decrease at the 45-minute (0.43) timepoint. This decrease in fluorescence then tapers off from 60 minutes (0.33) onwards with a return to baseline levels of STRIDE, akin to the negative control and ATMi only conditions. Conversely, γH2AX signal is at a minimum at the 0-minute timepoint (0.34), with increases at the 15- (0.40), 30- (0.54), and 45-minute (0.60) timepoints. This signal then plateaus at the 60-minute timepoint (0.58), and then reduces slightly as activation persists at 120- (0.52), 240- (0.53) and 1440-minutes (0.43) post-IR. Noticeably, although the fluorescence intensity of STRIDE decreases back to baseline levels, γH2AX is still active through to 1440-minutes.

**Figure 4:**
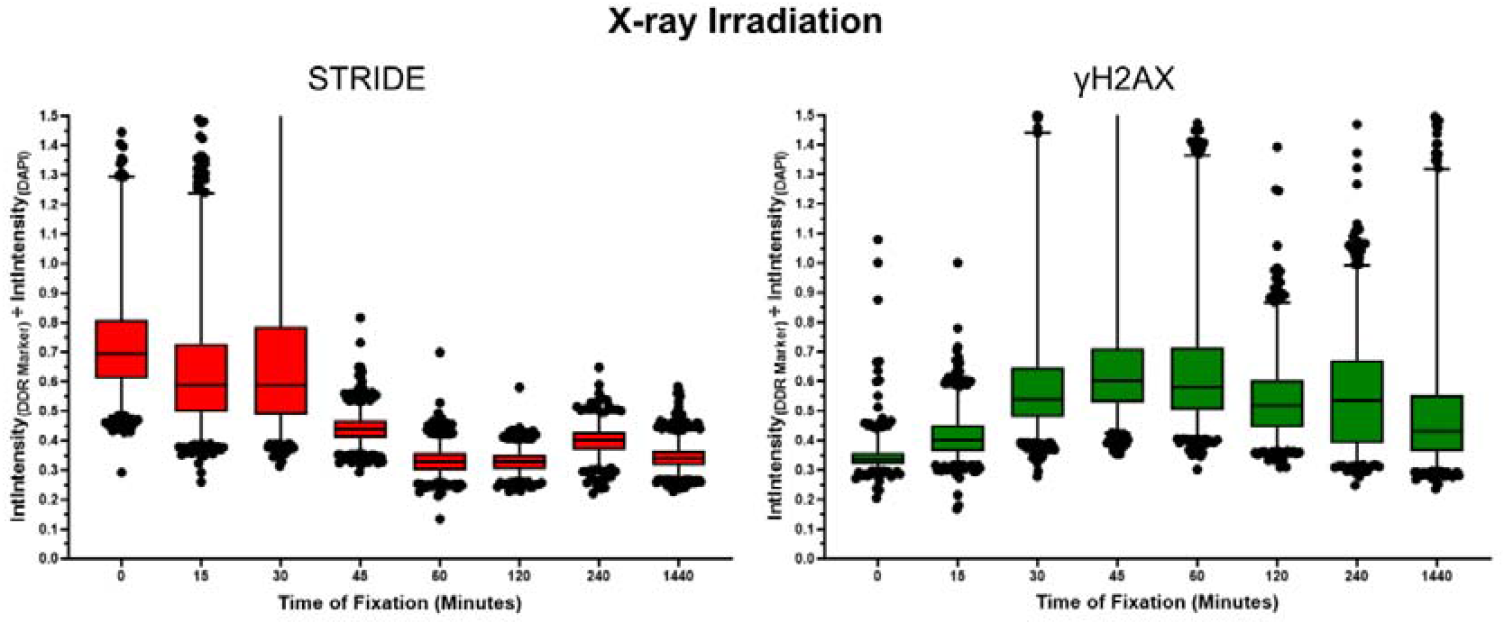
4T1 cells irradiated with X-rays. Integrated intensity values for STRIDE, γH2AX, and DAPI from single cells were measured. STRIDE and γH2AX values were normalised to DAPI. Data show the median value within the 25^th^ and 75^th^ percentile data points (box). Whiskers show the 2.5 - 97.5 percentile, and dots represent data points outside the range represented by whiskers.

One trend that is immediately noticeable is the characteristic ‘crossing over’ of integrated intensity between STRIDE and γH2AX at the 30- and 45-minute timepoints. As repair signalling increases, represented by γH2AX, STRIDE intensity decreases as a function of time beginning at 30 minutes and continuing through to the 1440-minute timepoint. Conversely, γH2AX increases from the 30-minute timepoint and then reaches a peak at both 45- and 60-minutes post-IR. This suggests that initiation of DNA damage repair obscures the ability of STRIDE to detect DSBs, likely due to recruitment of proteins to DSB ends that prevent STRIDE from binding.

This study also investigated in whether there was a cell phase dependence on STRIDE vs γH2AX normalised intensities, aiming to determine whether DNA damage or repair are enhanced within certain subpopulations. Specifically, how radiosensitive (G2/M) versus more radioresistant (G1 and S) phases respond to X-rays. By analysing the integrated intensities of DAPI and segmenting cell cycle phase, it appears that γH2AX is first active in G2/M phase cells at 30- and 45-minutes, with G1 cells lagging with elevated signals from 60-minutes onwards (Figure S2 to Figure S9). S phase cells remained relatively unchanged in terms of damage or repair accumulation. However, confocal microscopy provides limited fidelity in assessing DNA content across the cell cycle phase and these trends require high-throughput validation via flow cytometry.

Similarly to the AZD1390 only condition, inhibition of ATM with AZD1390 results in a low level of γH2AX phosphorylation across all timepoints (0.26 - 0.41), even when in combination with X-ray irradiation (Figure 5). This validates that even in the presence of radiation induced DSBs, that inhibition of ATM biological repair proteins will suppress the repair response. However, compared to irradiation alone, STRIDE intensities remain at their peak intensities at 0- (0.84), 15- (0.82) and 30-minutes (0.89) for a longer duration, and consequently decrease at later timepoints. Interestingly, the intensity of STRIDE signal across these timepoints is higher than with X-ray irradiation alone (Mann-Whitney p < 0.0001).

**Figure 5:**
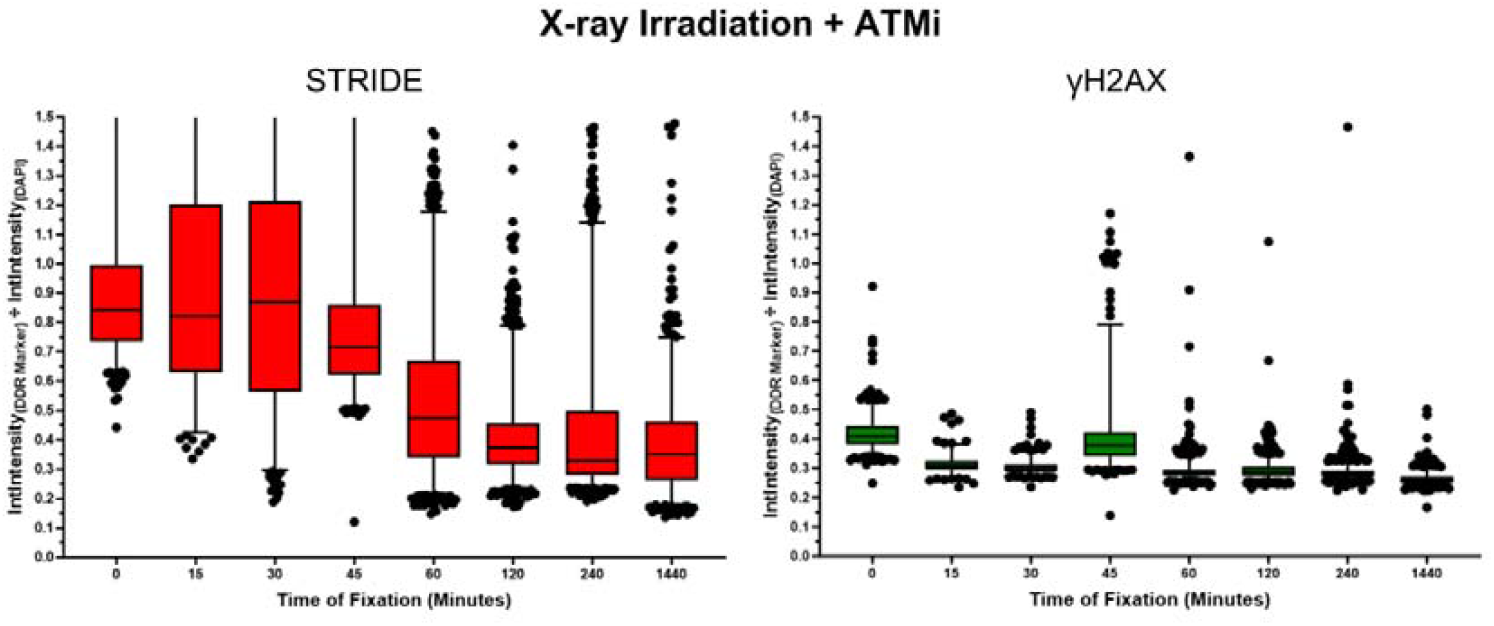
4T1 cells irradiated with X-rays and pre-treated with 30 nM AZD1390. Integrated intensity values for STRIDE, γH2AX, and DAPI from single cells were measured. STRIDE and γH2AX values were normalised to DAPI. Data show the median value within the 25^th^ and 75^th^ percentile data points (box). Whiskers show the 2.5 - 97.5 percentile, and dots represent data points outside the range represented by whiskers.

STRIDE signal then begins decreasing at 45- (0.72) and more substantially at 60-minutes (0.48) post-IR. This is also distinct from X-ray irradiation only, where STRIDE signal decreases substantially at the 45-minute (0.43) and 60-minute (0.33) timepoints, compared to when AZD1390 is added (p < 0.0001). One potential explanation for this delay in DNA damage repair is that upon addition of AZD1390, suppression of both ATM and γH2AX phosphorylation promotes alternate damage signalling molecules to compensate and commence accumulation of DDR proteins at DSB sites. These processes are likely less efficient than ATM or γH2AX, causing DSB repair proteins to take longer to accumulate at DSB sites, which delays their ability to obscure STRIDE detection.

### Colocalization of STRIDE and γH2AX Foci

To further investigate the relationship between STRIDE and γH2AX, the spatiotemporal correlation between the two markers was then explored. Specifically, in determining whether a γH2AX focus appears at the same location as a STRIDE focus when it resolves. Literature indicates that γH2AX foci can form 200 – 400 kilobases away from the DSB site [27, 28], so assessing colocalization could determine if γH2AX foci align with individual DSBs. Whether the disappearance of a STRIDE focus coincides with the emergence of a γH2AX focus, or rather if DSB resolution and the initiation of γH2AX repair signalling are linked, was also investigated (Figure 6).

**Figure 6:**
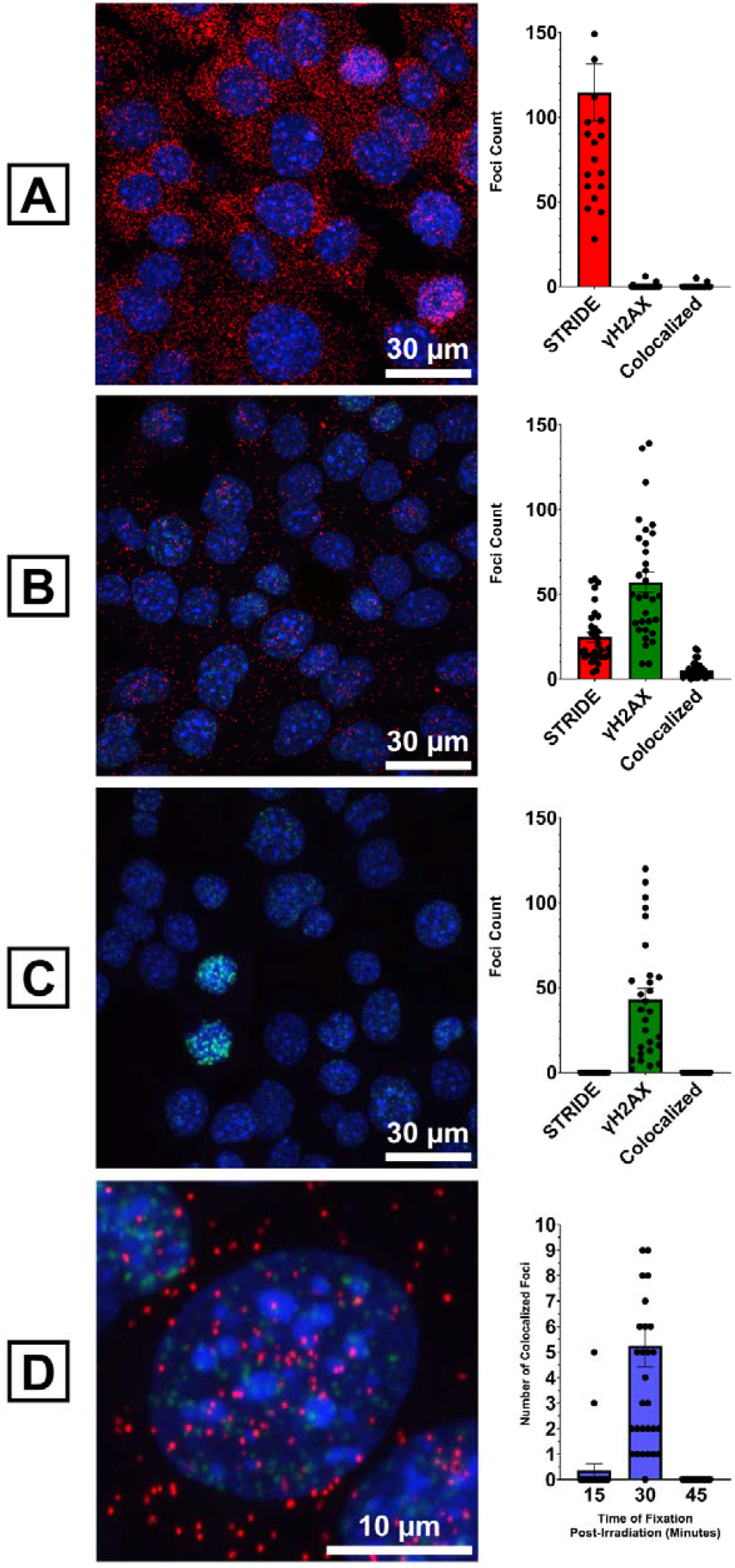
63x magnification images of X-ray irradiated 4T1 cells stained with DAPI (nuclei), STRIDE (red) and anti-γH2AX Alexa 488 conjugate (green), as well as corresponding foci counts at A.) 15-minutes post-IR, B.) 30-minutes post-IR, and C.) 45-minutes post-IR. Each image is representative of an independent well. D.) 30-minutes post-IR focusing on a single 4T1 cell. Mean + standard error of the mean values of colocalized foci identified across all three fixation timepoints.

From the images depicted in Figure 6, STRIDE and γH2AX foci generally do not colocalize, indicating that there may be proteins or other factors preventing STRIDE from binding to DSBs after their formation but before γH2AX foci are established (via H2AX phosphorylation). This is important as it identifies that the two markers represent distinctly different parts of the DSB formation and response pathways and are directly complementary. At the 30-minute timepoint there is a strong co-existence of STRIDE and γH2AX foci as STRIDE foci counts decrease, γH2AX counts increase. To confirm that the colocalization of foci at 30-minutes was not an artifact of the maximum intensity projection, individual z-slices were qualitatively analysed, showing fewer instances of colocalization across z-slices than with the maximum projection. This is indicative that that STRIDE and γH2AX markers do not frequently colocalize on the same z-plane.

## Conclusion

This study supports the critical role of ATM in the repair of DSBs and highlights the potential of AZD1390 to enhance the radiosensitivity of cancerous cells. By employing STRIDE, the effect of AZD1390 on DNA repair kinetics within a 24-hour timeframe in 4T1 cells was successfully characterised. It was discovered that the physical presence of DNA breaks can still be detected using STRIDE when cells are treated with AZD1390, making it a superior tool for assessing DNA damage formation in scenarios where DDR markers like γH2AX are suppressed, and are hence unreliable indicators of genotoxicity. Moreover, STRIDE showed that treatment of cells with a combination of AZD1390 and X-rays delayed the onset of DSB resolution compared to X-rays alone. Obfuscation of the DSB to STRIDE due to DSB sensing is delayed in the presence of ATMi, indicating ATMis play a greater role in suppressing DDR beyond ATM inhibition alone. Consequently, ATM inhibition effectively prolongs the persistence of radiation-induced DNA damage, providing a potential therapeutic advantage by stalling DNA sensing and repair in radioresistant cell types and thereby extending the window during which they remain vulnerable to radiotherapy. In future studies, testing of AZD1390 against multiple cancer cell types with varying radioresistance would provide valuable insights into whether its effectiveness in perturbing DSB repair is universal. Exploring combination therapies that pair AZD1390 with other DDR-targeting chemotherapeutics could also provide a more comprehensive strategy to both overcome radioresistance and enhance the DSB burden in aggressive or late-stage cancers.

## Acknowledgements

This research was funded by the Tour de Cure PhD Scholarship, grant ID number RSP-641-FY2023. We acknowledge Tour de Cure’s contribution towards the preparation of this manuscript through the Tour de Cure Scholarship Grant and appreciate their continued support for this work.

We acknowledge Microscopy Australia (ROR: 042mm0k03) at the Future Industries Institute, University of South Australia, enabled by NCRIS. Thanks to Dr. Alex Cavallaro for assistance with operation of the Zeiss LSM 710 confocal microscope.

Many thanks to Dr. Bill Liapis for assisting with the operation of the X-ray irradiator.

## Supplemental Information

Document S1. Figures S1-S10 and Table S1.

