## Supplementary Data for "Mechanisms of ATM Inhibitor AZD1390-Mediated Radiosensitization by Comparing DNA DSB Formation and Repair in 4T1 Cells"

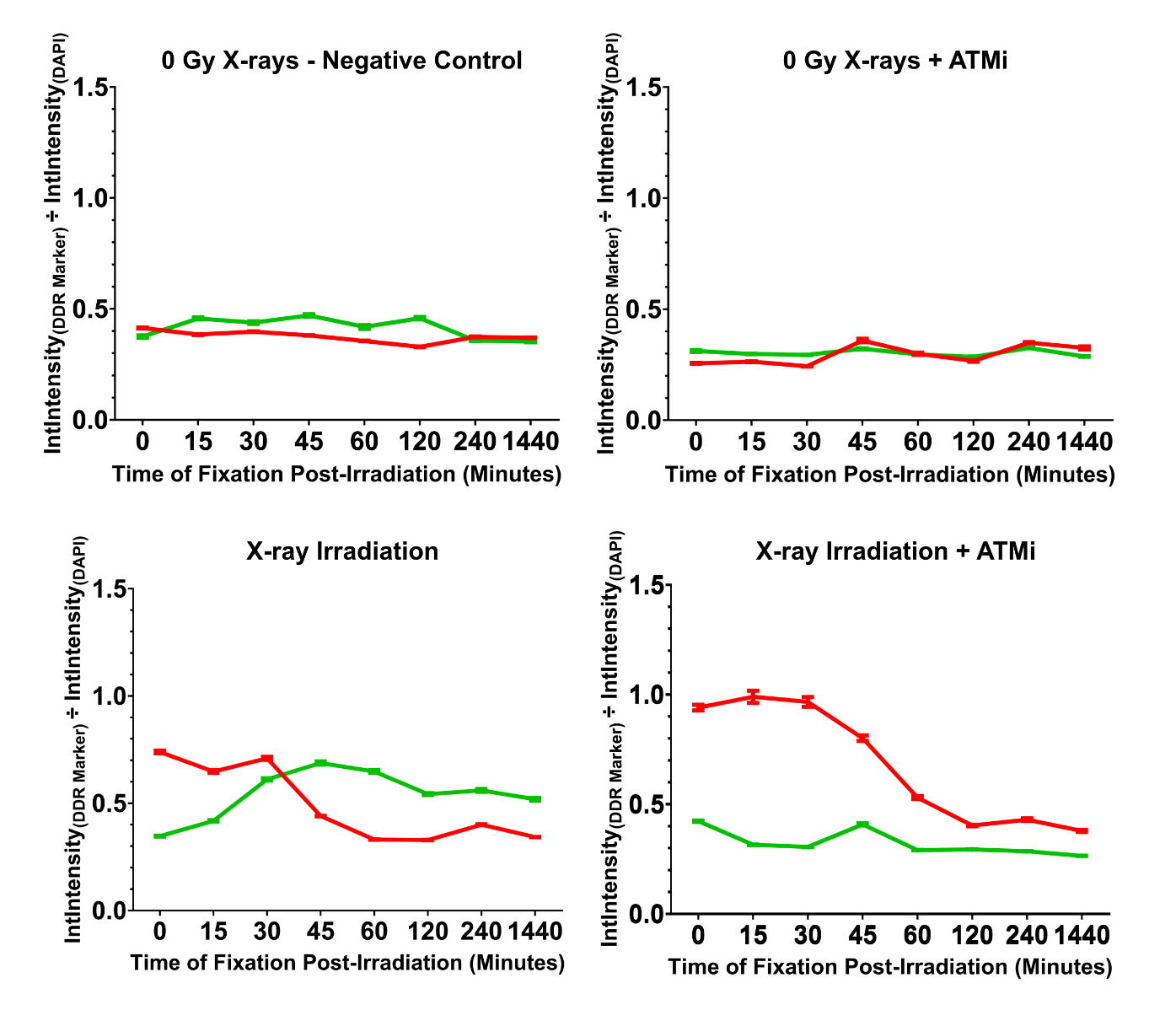

Figure S1: Mean and standard error of the mean (SEM) values of integrated intensity for STRIDE (red) and γH2AX (green) normalized to DAPI for 4T1 cells under various conditions. Top left: no X-ray irradiation or pre-treatment with AZD1390, top right: exclusively pre-treatment with 30 nM AZD1390, bottom left: exclusively X-ray irradiation, bottom right: X-ray irradiation in combination with 30 nM AZD1390 pre-treatment. Integrated intensity measurements were taken from individual nuclei across different fixation timepoints.

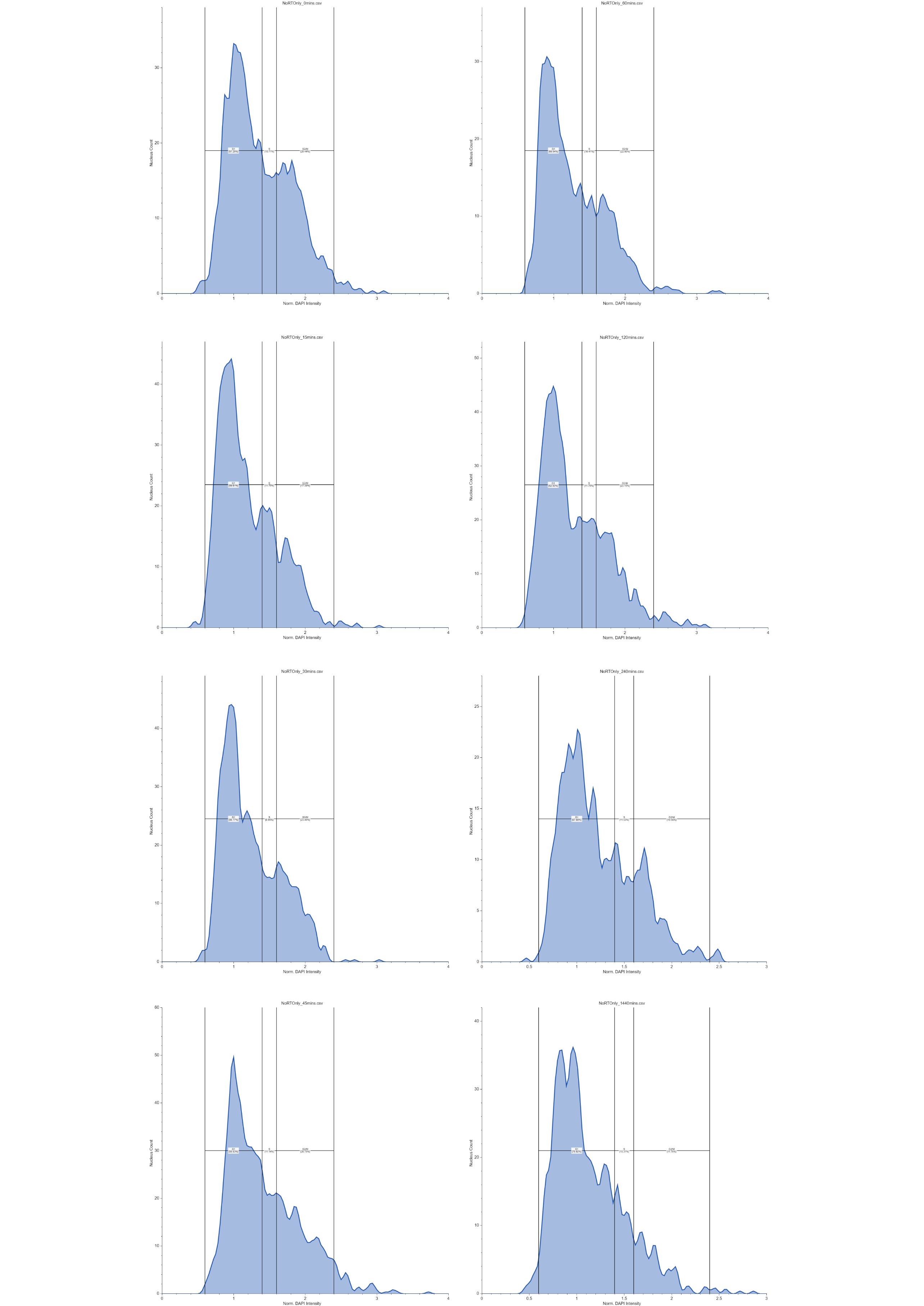

Figure S2: No X-rays or AZD1390. Histograms depicting the integrated intensity of DAPI normalized to the intensity of the 2N peak for each fixation timepoint. The gating of the G1 (0.6 – 1.4), S (1.4 – 1.6), and G2/M (1.6 – 2.4) phases are also displayed. Plots produced with Floreada.io.

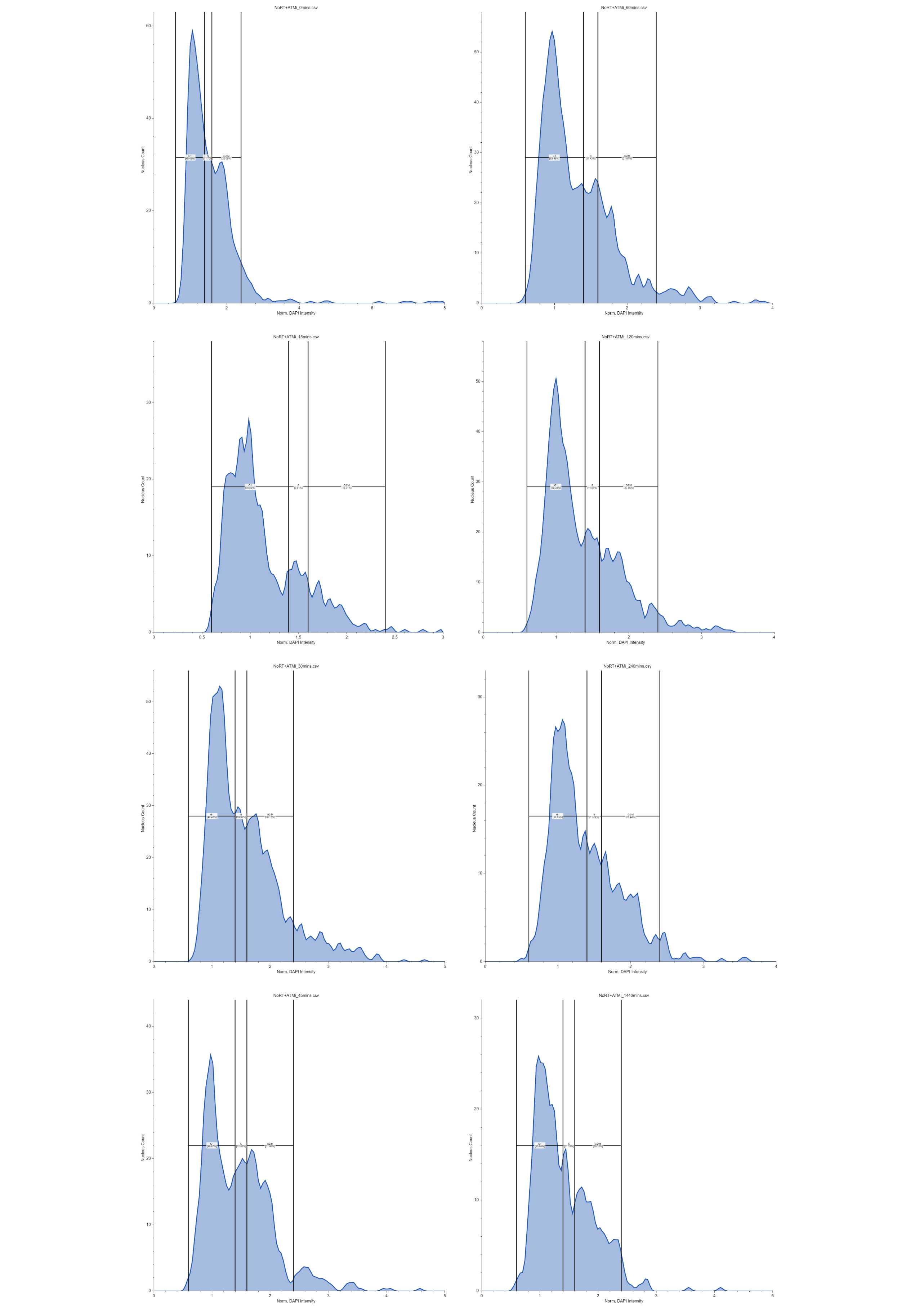

Figure S3: No X-rays + AZD1390. Histograms depicting the integrated intensity of DAPI normalized to the intensity of the 2N peak for each fixation timepoint. The gating of the G1 (0.6 – 1.4), S (1.4 – 1.6), and G2/M (1.6 – 2.4) phases are also displayed. Plots produced with Floreada.io.

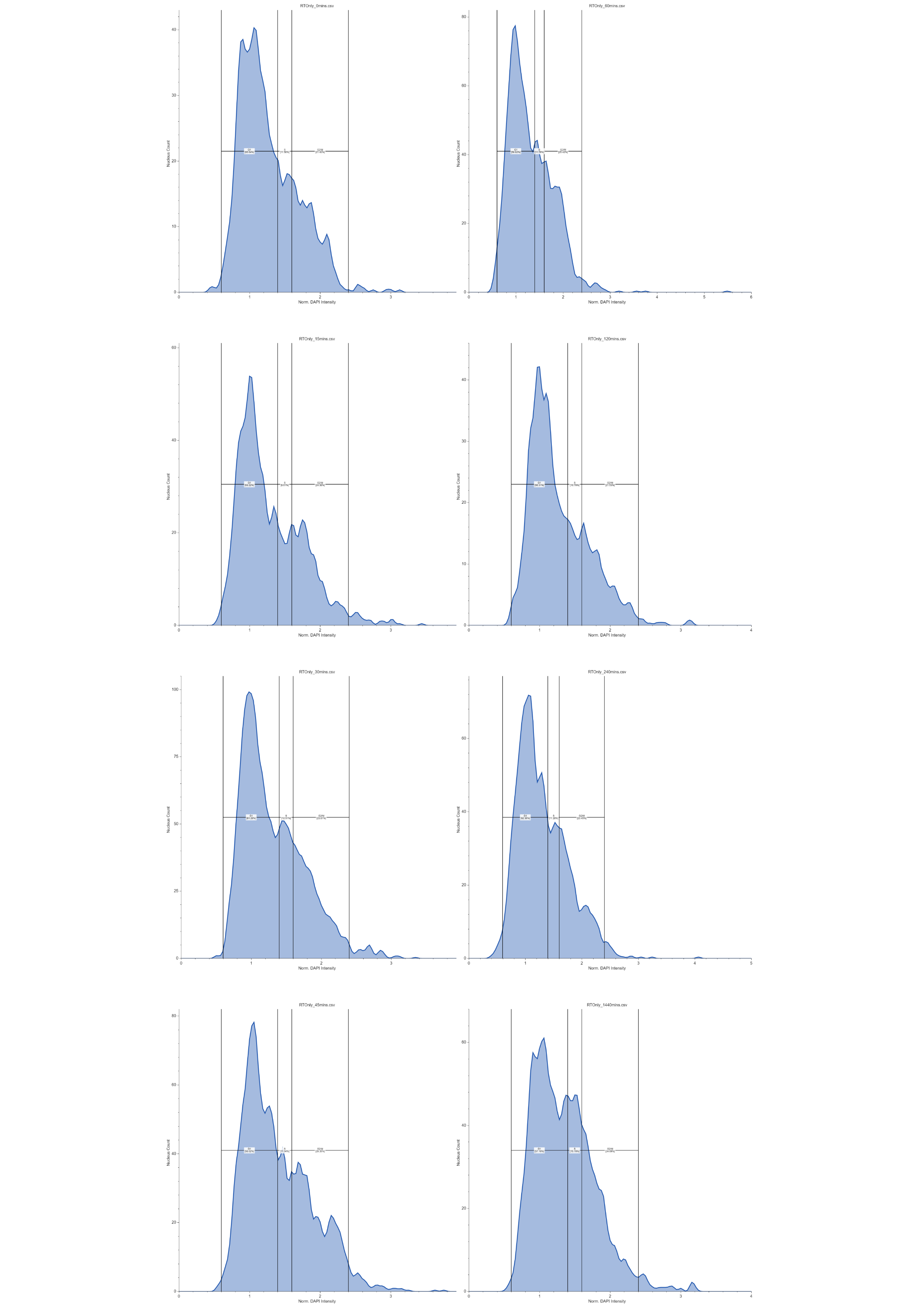

Figure S4: X-ray irradiation only. Histograms depicting the integrated intensity of DAPI normalized to the intensity of the 2N peak for each fixation timepoint. The gating of the G1 (0.6 – 1.4), S (1.4 – 1.6), and G2/M (1.6 – 2.4) phases are also displayed. Plots produced with Floreada.io.

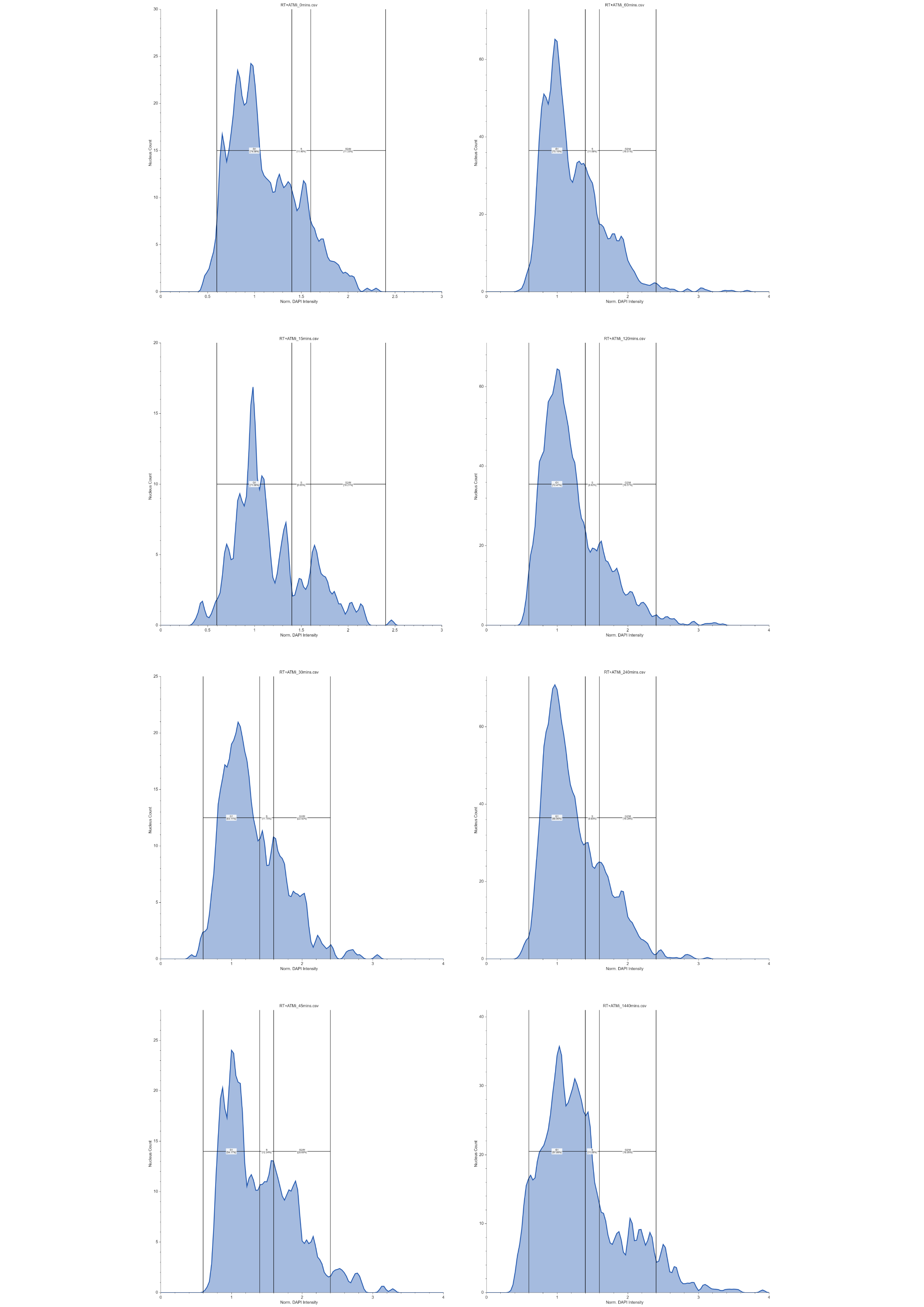

Figure S5: X-ray irradiation + AZD1390. Histograms depicting the integrated intensity of DAPI normalized to the intensity of the 2N peak for each fixation timepoint. The gating of G1 (0.6 – 1.4), S (1.4 – 1.6), and G2/M (1.6 – 2.4) phases are also displayed. Plots produced with Floreada.io.

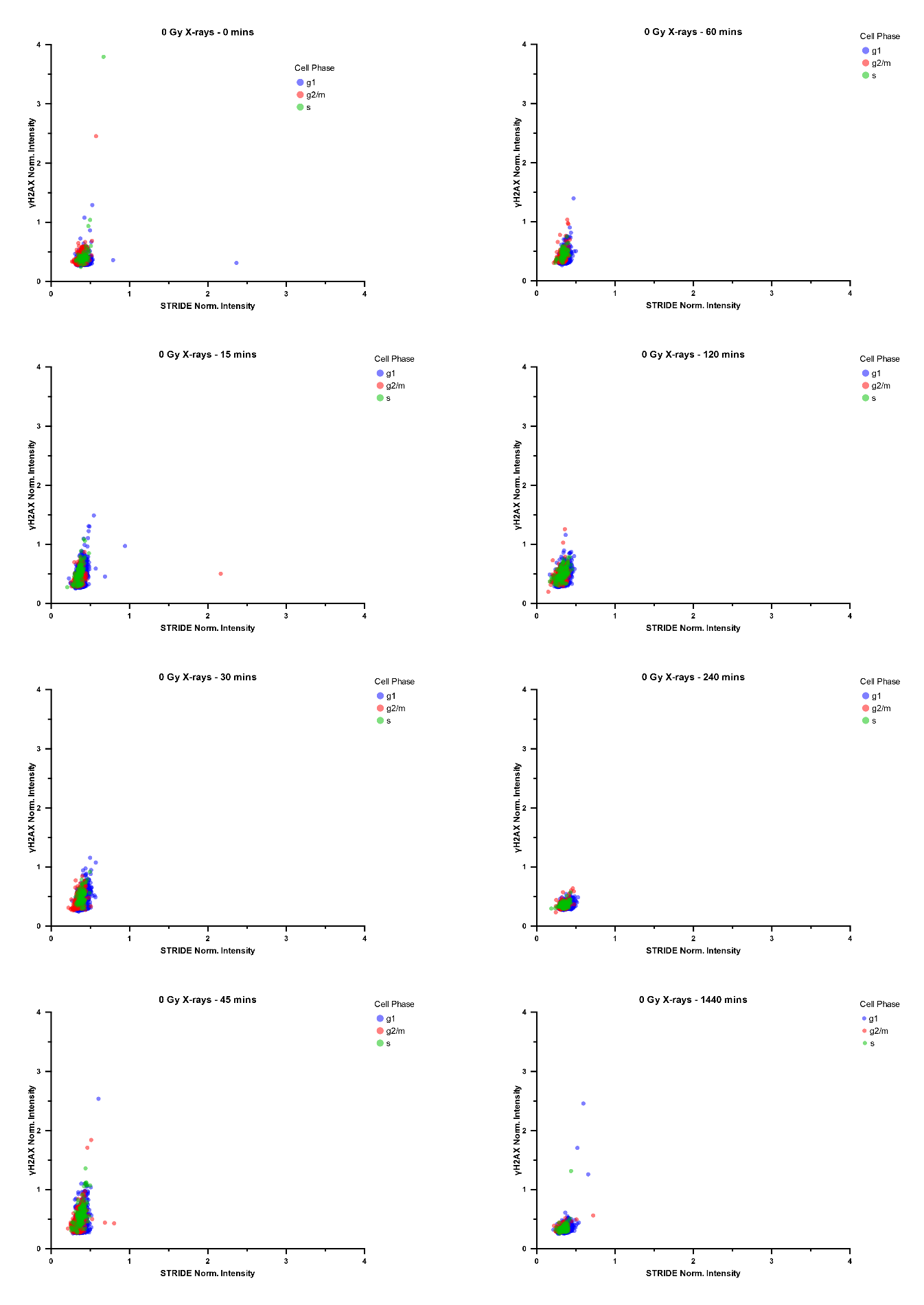

Figure S6: Normalized integrated intensities of STRIDE (x-axis) versus γH2AX (y-axis) for 4T1 cells with no X-ray or AZD1390 treatment. Intensities were gated (Figure S2) to determine the approximate proportion of cells in G1 (blue), S (green) and G2/M (red) phases.

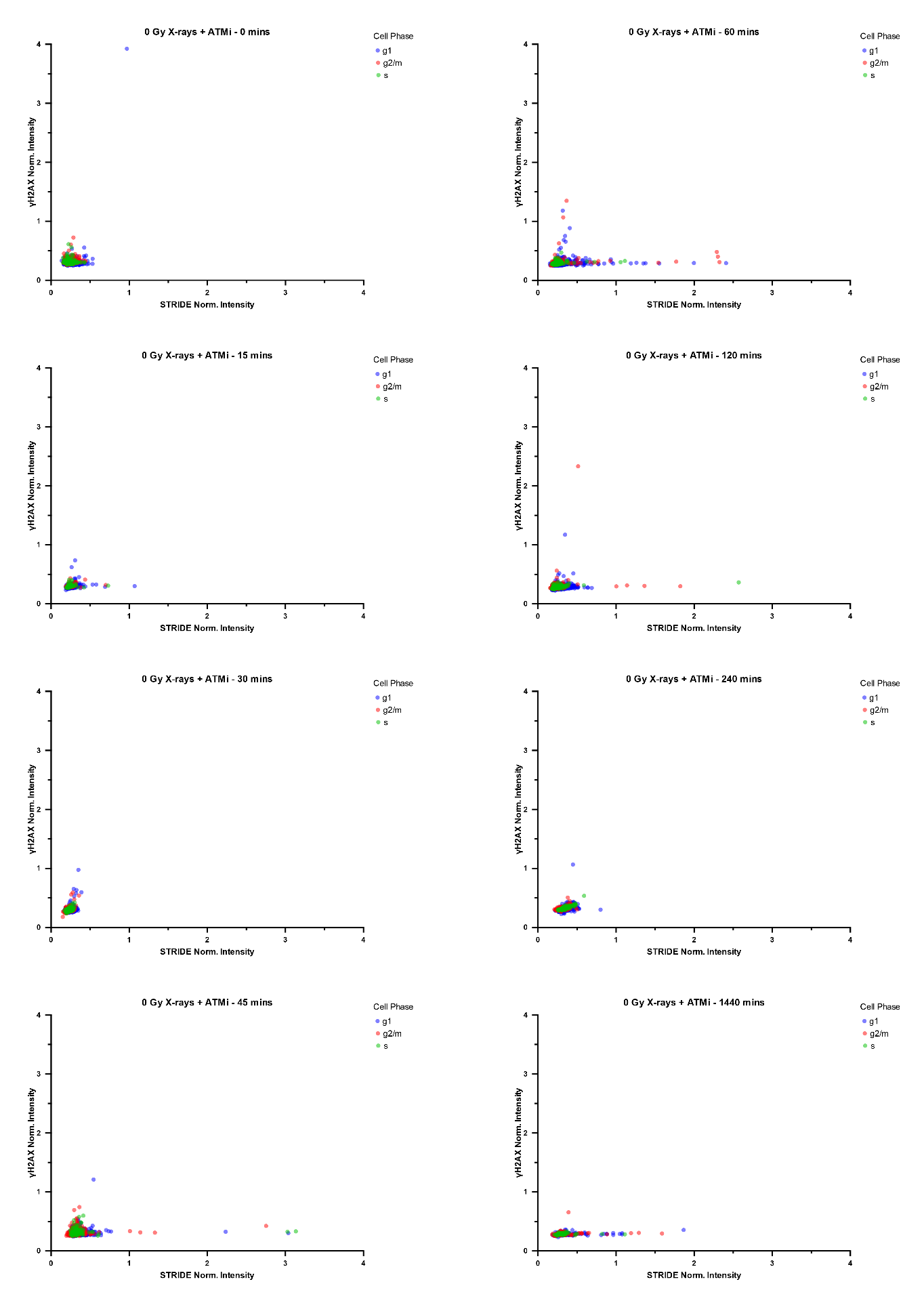

Figure S7: Normalized integrated intensities of STRIDE (x-axis) versus γH2AX (y-axis) for 4T1 cells pre-treated with 30 nM AZD1390 only. Intensities were gated (Figure S3) to determine the approximate proportion of cells in G1 (blue), S (green) and G2/M (red) phases.

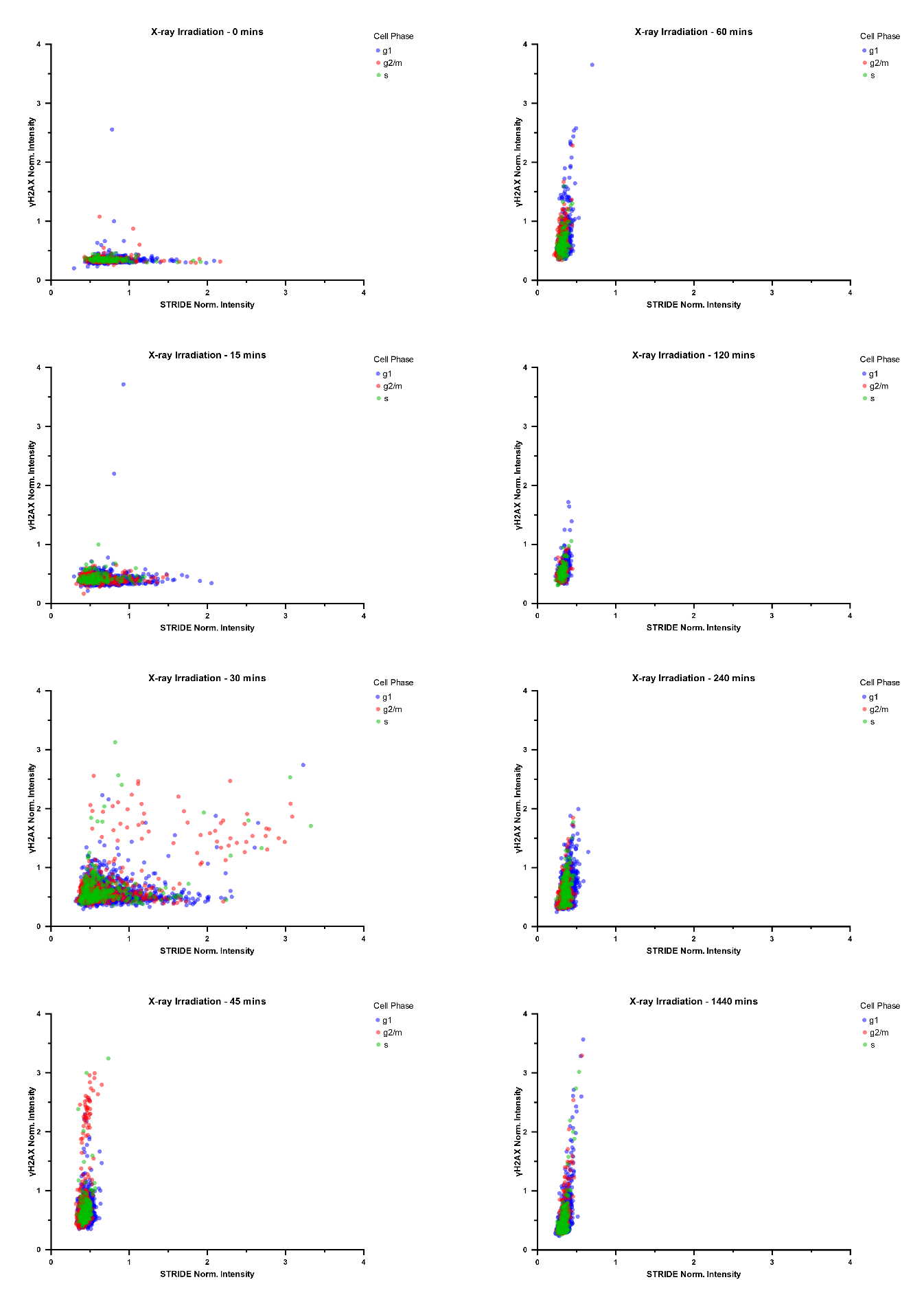

Figure S8: Normalized integrated intensities of STRIDE (x-axis) versus γH2AX (y-axis) for 4T1 cells treated with X-rays only. Intensities were gated (Figure S4) to determine the approximate proportion of cells in G1 (blue), S (green) and G2/M (red) phases.

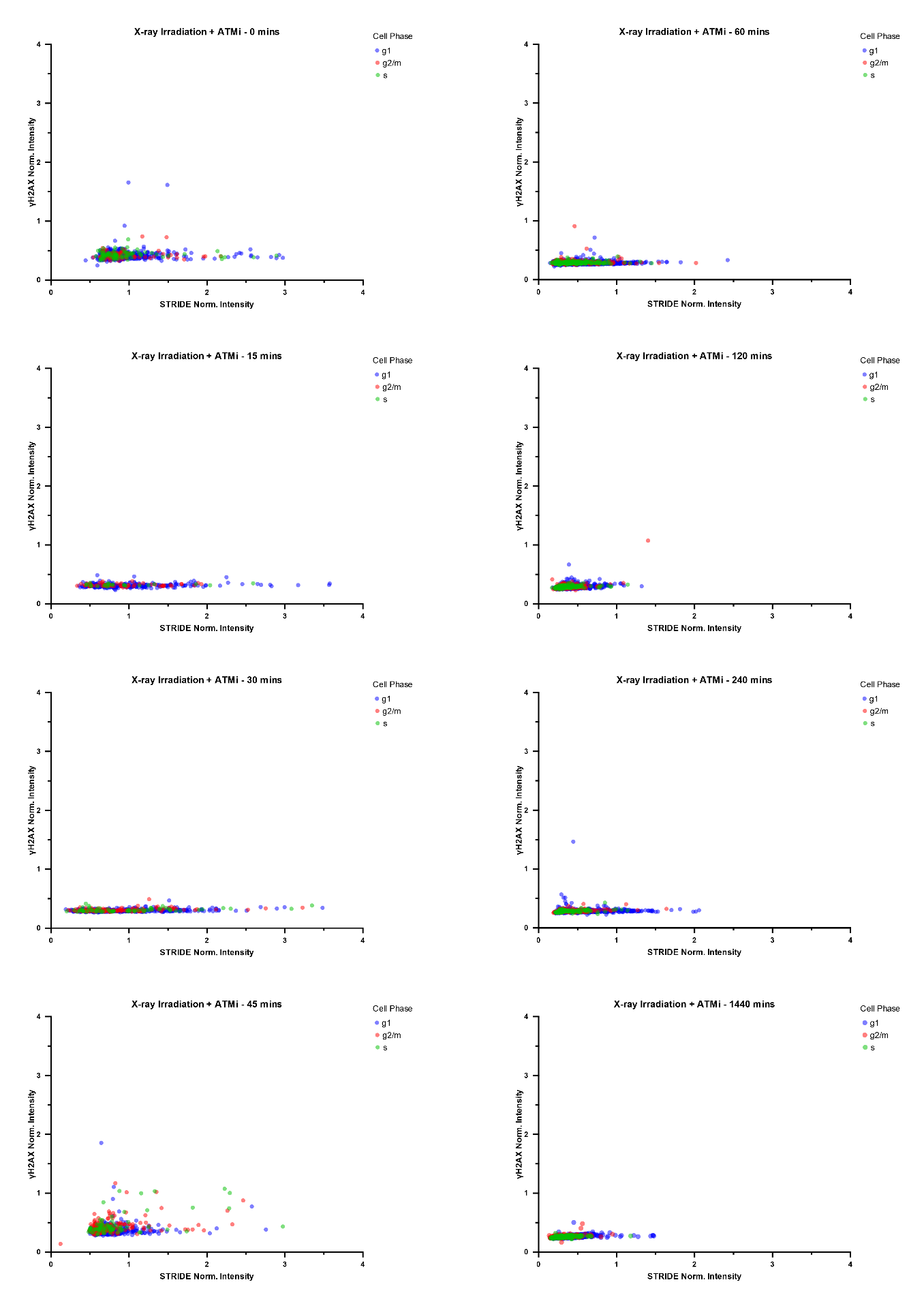

Figure S9: Normalized integrated intensities of STRIDE (x-axis) versus γH2AX (y-axis) for 4T1 cells treated with both X-rays and 30 nM AZD1390. Intensities were gated (Figure S5) to determine the approximate proportion of cells in G1 (blue), S (green) and G2/M (red) phases.

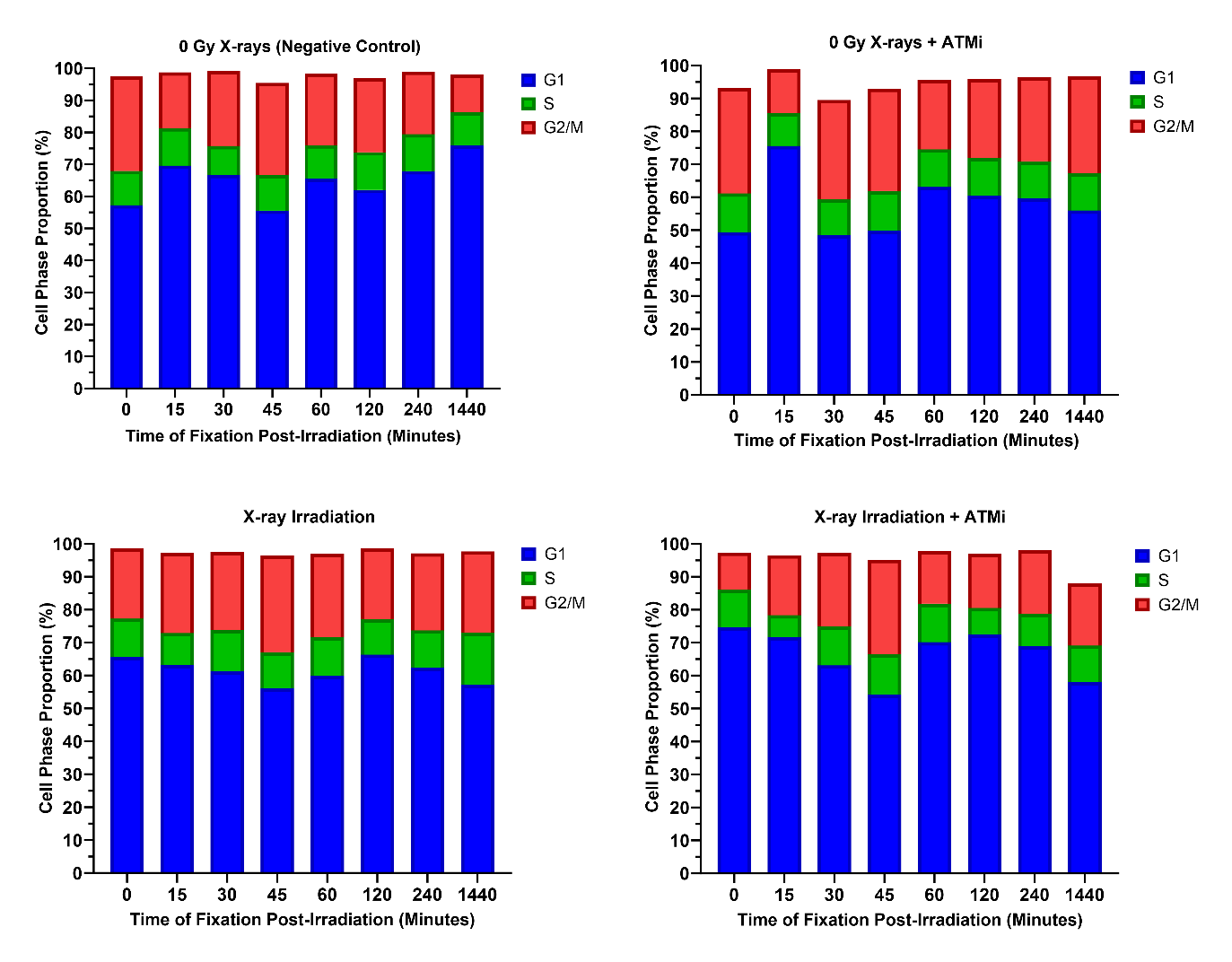

Figure S10: Stacked box plots depicting the percentage of 4T1 cells in each cell phase, G1 (blue), S (green), and G2/M (red) across fixation timepoints post-IR. Cells were gated based on the ranges defined in Figure S2-S5.

Table S1: Descriptive statistics for each dataset for STRIDE and H2AX across all timepoints. Values recorded include the number of values in each population (n), mean, median, standard deviation, standard error of the mean, range (defined by Prism as the difference in the minimum and maximum values), and 25th and 75th percentiles.

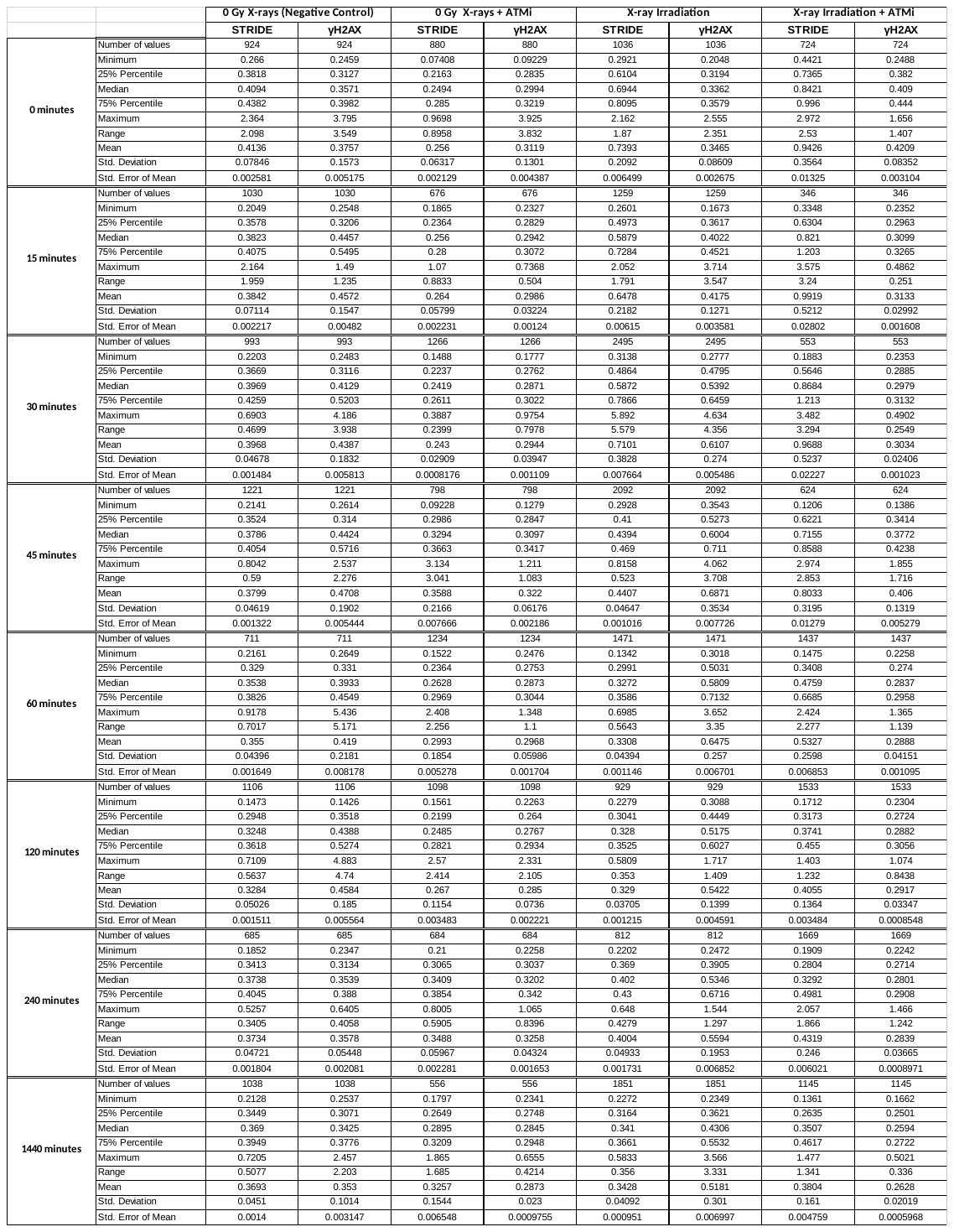
